# Removal of maternal compensation reveals retinal-specific roles for Pomgnt1 in zebrafish

**DOI:** 10.64898/2026.11.02.763738

**Authors:** Callista D. Oliveira, Kyle P. Flannery, Lillian M. Simhon, Nuo Shi, Deze Liu, Daniel Burbano, M. Chiara Manzini

## Abstract

Genetic variants in *Protein O-mannose N-acetylglucosaminyltransferase 1* (*POMGNT1*) result in a broad spectrum of disorders, ranging from severe congenital muscular dystrophy with brain and ocular malformations to non-syndromic retinitis pigmentosa. Despite POMGNT1’s established role in the glycosylation of the transmembrane glycoprotein α-dystroglycan (α-DG), the mechanistic basis for this clinical variability remains poorly understood. Previous work in a zebrafish *pomngt1* knock-out (KO) line identified retinal degeneration at 6 months post-fertilization (mpf). However, potential involvement of other tissues was not characterized, and compensation from maternal transcripts which can occur in the zebrafish oocyte was not tested. Here, we analyzed maternal-zygotic *pomgnt1* KOs (MZKOs) to determine whether maternal compensation influences the onset and severity of *pomgnt1*-associated phenotypes. We assessed survival, gross morphology, locomotor activity, behavioral patterns, and retinal organization across multiple timepoints. MZKOs survived to adulthood and exhibited no morphological or locomotor deficits up to 5 mpf. Retinal abnormalities were found as early as 2 mpf in MZKOs, demonstrating that maternally provided factors of *pomgnt1* delay the onset of retinal pathology. Additionally, MZKOs displayed a novel “wall-hugging” behavioral pattern that may be consistent with visual impairment or anxiety. These findings demonstrate a strong tissue-specific phenotype following loss of *pomgnt1* in zebrafish showing evolutionary divergence in the necessity for Pomgnt1 function in the muscle. However, this strain provides a useful system for investigating the mechanisms underlying the retinal vulnerability observed in POMGNT1-related retinitis pigmentosa.

## Introduction

Protein O-mannose N-acetylglucosaminyltransferase 1 (POMGNT1) is a glycosyltransferase enzyme mutated in rare autosomal recessive disorders ranging from severe Congenital Muscular Dystrophy (CMD) with brain involvement to non-syndromic retinitis pigmentosa (RP) (Walimbe et al., 2026; Xu et al., 2016). The severe CMD presentation associated with POMGNT1 loss-of-function is also termed Muscle Eye Brain disease (MEB), and it includes brain malformations (simplified gyral pattern or pachygyria, enlarged ventricles, and cerebellar hypoplasia) and ocular deficits such as progressive myopia, retinal degeneration, glaucoma, and cataracts (Hehr et al., 2007; Yoshida et al., 2001). However, *POMGNT1* variants can also result in CMD with intellectual disability or later onset limb girdle muscular dystrophy (LGMD) without cognitive deficits (Clement et al., 2008). In parallel, presentations with no muscle disease have been identified in multiple cases of non-syndromic RP and retinal degeneration (Patel et al., 2023; Xu et al., 2016; Ziccardi et al., 2025).

Variable glycosylation efficiency has been suggested as the cause of differential susceptibility of multiple tissues in MEB versus the isolated retinal involvement in RP (Manya and Endo, 2017). POMGNT1 is involved in O-linked glycosylation of *α*-dystroglycan (*α*-DG), the extracellular glycoprotein of the Dystrophin-Glycoprotein Complex (DGC). Glycans on *α*-DG bind ligands in the extracellular matrix (ECM), forming a structural and signaling bridge between the intracellular actin cytoskeleton and extracellular environment (Imae et al., 2026; Kanagawa, 2021; Walimbe et al., 2026). These glycan interactions are essential in morphogenesis and maintenance of multiple tissues, including the skeletal muscle, brain, and retina (Hord et al., 2025; Jahncke and Wright, 2023; Kanagawa, 2021). *α*-DG-ECM interactions are primarily mediated through a long functional glycan, termed matriglycan, initiated by a heterocomplex of Protein O-mannosyltransferases 1 and 2 (POMT1/POMT2) and elongated by Protein O-mannose N-Acetylglucosaminyltransferase 2 (POMGNT2) (Halmo et al., 2017; Inamori et al., 2012, Manya et al., 2004). While POMGNT1 elongates different glycans initiated by the POMT complex, it has been found to partially contribute to matriglycan maturation (Imae et al., 2026; Kuwabara et al., 2016; Sheikh et al., 2017). However, the tissue-specific consequences of *POMGNT1* loss are not fully explained by its currently understood role in glycan synthesis.

A similar phenotypic discrepancy has been found in animal models of *POMGNT1* loss of function. *Pomgnt1* knock-out (KO) mice recapitulate the human presentation with muscle disease, neuronal migration deficits, and retinal abnormalities (Hu et al., 2010; Liu et al., 2006). In contrast, the only reported phenotype in *pomgnt1* zebrafish KOs is retinal degeneration at 6 months post-fertilization (mpf) (Liu et al., 2020). Loss of other *α*-DG glycosyltransferases (*pomt1*, *pomt2*, and *pomgnt2*) had shown unexpectedly mild phenotypes in standard zygotic zebrafish KOs (ZKO) obtained from heterozygous crosses (Flannery et al., 2026; Karas et al., 2024; Liu et al., 2022). As zebrafish are externally fertilized and can undergo maternal compensation, where maternally deposited factors (mRNA and proteins) buffer early developmental phenotypes, phenotypes in zygotic mutant models can underestimate the consequence of complete gene loss due to preserved maternal mRNA (Pelegri, 2003; Zhao et al., 2017). Full phenotypic presentation in *pomt1* and *pomgnt2* mutants could only be achieved in maternal-zygotic knockouts (MZKOs) obtained from ZKO mothers, which exhibited reduced survival, severe motor deficits, disrupted axon guidance, and retinal disorganization (Flannery et al., 2026; Karas et al., 2024). As *pomgnt1* mRNA is also maternally provided (Baia Amaral et al., 2024; White et al., 2017), it is possible that similar maternal compensation may influence phenotype presentation in *pomgnt1* mutants as well.

Here, we report the analysis of *pomgnt1* MZKO animals from the same strain where ZKO analysis was conducted (Liu et al., 2020). We investigated whether maternal loss of *pomgnt1* alters the timing and severity of phenotypic presentation in zebrafish. In addition to retinal deficits, we assessed whether *pomgnt1* loss produces broader muscular or locomotor deficits. There were no signs of muscle disease as late as 5 mpf. However, we found that photoreceptor deficits consistent with those observed in ZKOs were present as early as 2 mpf in MZKO indicating the presence of maternal compensation. Our results reveal a selective retinal role for Pomgnt1 glycosylation in zebrafish.

## Results

### Morphological in pomgnt1 MZKO fish

Previous studies on *pomgnt1* loss of function in zebrafish exclusively utilized ZKO progeny, leaving potential contribution of maternal compensation in mutant *pomgnt1* phenotypes unexplored (Liu et al., 2020). We raised ZKO females from the same CRISPR/Cas9 edited line (*pomgnt1^Syn7^*) to generate MZKO and maternal heterozygotes (MHet). Previous studies on ZKO and MZKO models of *α*-DG glycosylation has shown that glycosylation was preserved in ZKO larvae up to 7 days post-fertilization (dpf) (Flannery et al., 2026; Karas et al., 2024). To test whether this was the case for the *pomgnt1* line, we performed whole mount immunofluorescence using the glyco-specific antibody IIHC6 which recognizes *α*-DG glycosylation. Analysis on 7 dpf larvae obtained from crosses to generate either ZKOs or MZKOs confirmed that *α*-DG glycosylation is still present in ZKOs at this age (**Fig. 1A**) but absent in MZKOs (**Fig. 1B**).

**Figure 1.**
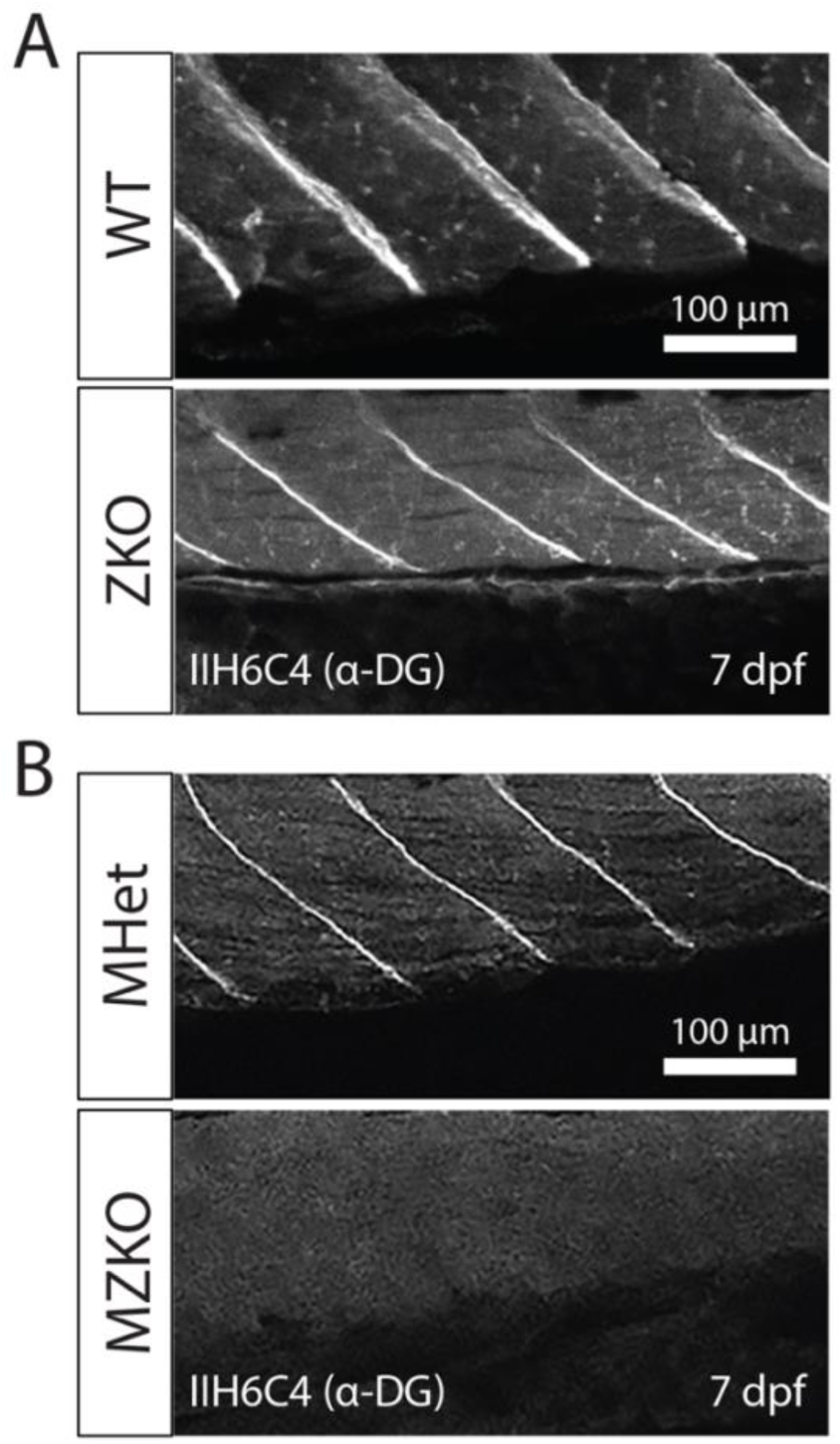
*α*-DG glycosylation following zygotic and maternal-zygotic loss of *pomgnt1*. **A.** Immunofluorescent whole mount images of tail muscle from WT and ZKO progeny using the IIH6 glyco-specific antibody demonstrate residual glycosylation of *α*-DG ZKOs at 7 dpf. **B.** The same experiment in MZKOs demonstrates the complete loss of *α*-DG glycosylation in MZKOs at 7 dpf. Scale bars: 100 µm

To determine whether the loss of *pomgnt1* affects the viability and gross morphology of MZKO zebrafish, we analyzed the survival rates and body length of MHets and MZKO animals across several developmental timepoints at 1, 2, and 5 mpf. We found that the MHet and MZKO progeny survived in the expected 1:1 Mendelian ratio across all timepoints, indicating no genotype-dependent differences in survival during initial growth and development and into early adulthood (1 mpf: MHet 49.50 ± 4.49%, n=56, MZKO 50.50 ± 4.49%, n=58; N=3 clutches; 2 mpf: MHet 49.70 ± 5.23%, n=75, MZKO 50.30 ± 5.23%, n=74; N=4 clutches; 5 mpf: MHet 42.67 ± 2.59%, n=27, MZKO 57.33 ± 2.59%, n=36; N=3 clutches) (**Fig. 2A**).

**Figure 2.**
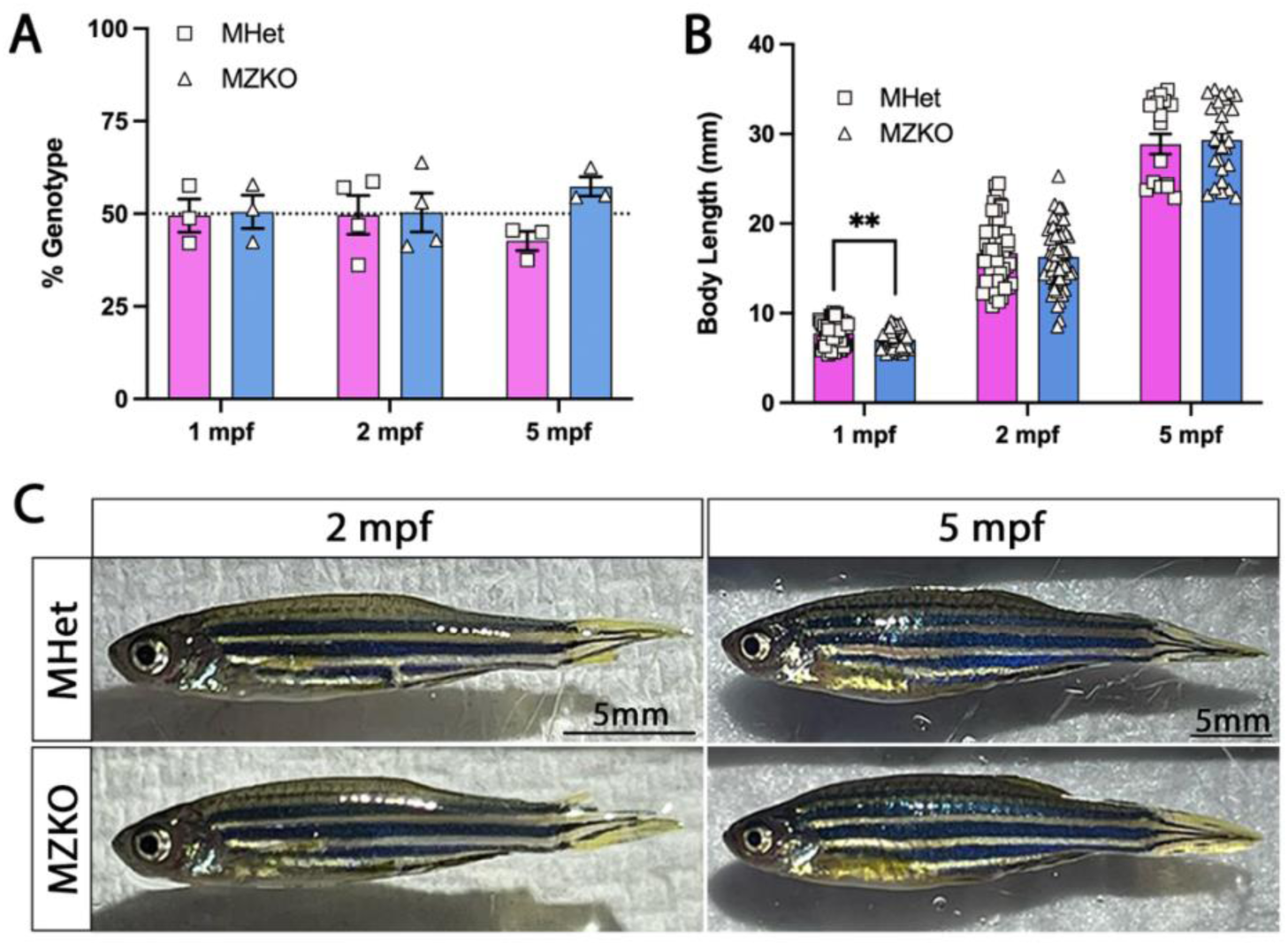
No gross phenotypes are noted following maternal-zygotic loss of *pomgnt1*. **A.** Survival rates at 1, 2, and 5 mpf reveal no impact of *pomgnt1* loss on MZKO overall viability compared to MHet controls. Datapoints indicate independent clutches. Data are presented as mean ± SEM. **B.** Average body length of MHet and MZKO fish across multiple cohorts at 1, 2, and 5 mpf reveal significant difference in growth at 1 mpf only, with pattern lost at 2 and 5 mpf. Datapoints indicate individual fish. **C.** Visual comparison of anatomy and coloration of MHet and MZKO fish at the 2mpf and 5mpf timepoints reveal no changes in gross body morphology into adulthood following *pomgnt1* loss. Scale bars: 5 mm

We evaluated gross body morphology by measuring body length and found statistically significant differences between MHets and MZKOs at 1 mpf (1 mpf: MHet 7.769 ± 0.19 mm, n=52, MZKO 7.047 ± 0.15 mm, n=50; N=3 clutches; p=0.0036), with this trend disappearing at later timepoints (2 mpf: MHet 16.71 ± 0.47 mm, n=55, MZKO 16.29 ± 0.45 mm, n=59; N=3; p=0.591; 5 mpf: MHet 28.87 ± 1.12 mm, n=18, MZKO 29.34 ± 0.85 mm, n=25; N=2; p=0.996) (**Fig. 2B**). Further visual evaluation of the zebrafish at the same timepoints revealed no differences in general morphology between MHets and MZKOs as they developed into adulthood (**Fig. 2C**). Overall, there was no reduction in lifespan or the presence of any other observable differences in body morphology in progeny lacking maternal compensation. The early difference in body length was not maintained at later timepoints, and further analysis is needed to determine whether growth trajectories differ between the two groups.

### pomgnt1 MZKO fish do not show locomotor deficits but have altered swimming patterns

Although maternal zygotic *pomgnt1* loss did not demonstrate any negative impacts on survival or gross morphology, dystroglycanopathies are primarily characterized by progressive muscle weakness and degeneration. To test whether swimming ability was affected, we measured multiple parameters of locomotor activity via automated video tracking in circular arenas across developmental timepoints in MHets and MZKOs. At 1 mpf, no significant differences were found across swim distance (MHet: 682.54 ± 28.98 cm, n=37; MZKO: 700.47 ± 33.93 cm, n=44; N=2; p=0.695), maximum velocity (MHet: 11.88 ± 0.75 cm/s, n=37; MZKO: 12.87 ± 0.29 cm/s, n=44; N=2; p=0.4616), or maximum acceleration (MHet: 267.95 ± 18.08 cm/s^2^, n=37; MZKO: 304.88 ± 14.91 cm/s^2^, n=44; N=2; p=0.116) (**Fig. 3A-C**). The same preservation of locomotor activity was found at the 5 mpf timepoint across distance (MHet: 3843.4 ± 318.06 cm, n=18; MZKO: 4374.0 ± 295.38 cm, n=22; N=2; p=0.2303), maximum velocity (MHet: 8.32 ± 0.68 cm/s, n=18; MZKO: 9.73 ± 0.69 cm/s, n=22; N=2; p=0.1553), and maximum acceleration (MHet: 29.18 ± 3.35 cm/s^2^, n=18; MZKO: 30.86 ± 2.66 cm/s^2^, n=22, N=2; p=0.6940) (**Fig. 3D-F**). These results indicate that loss of *pomgnt1* does not significantly impair gross locomotor function.

**Figure 3.**
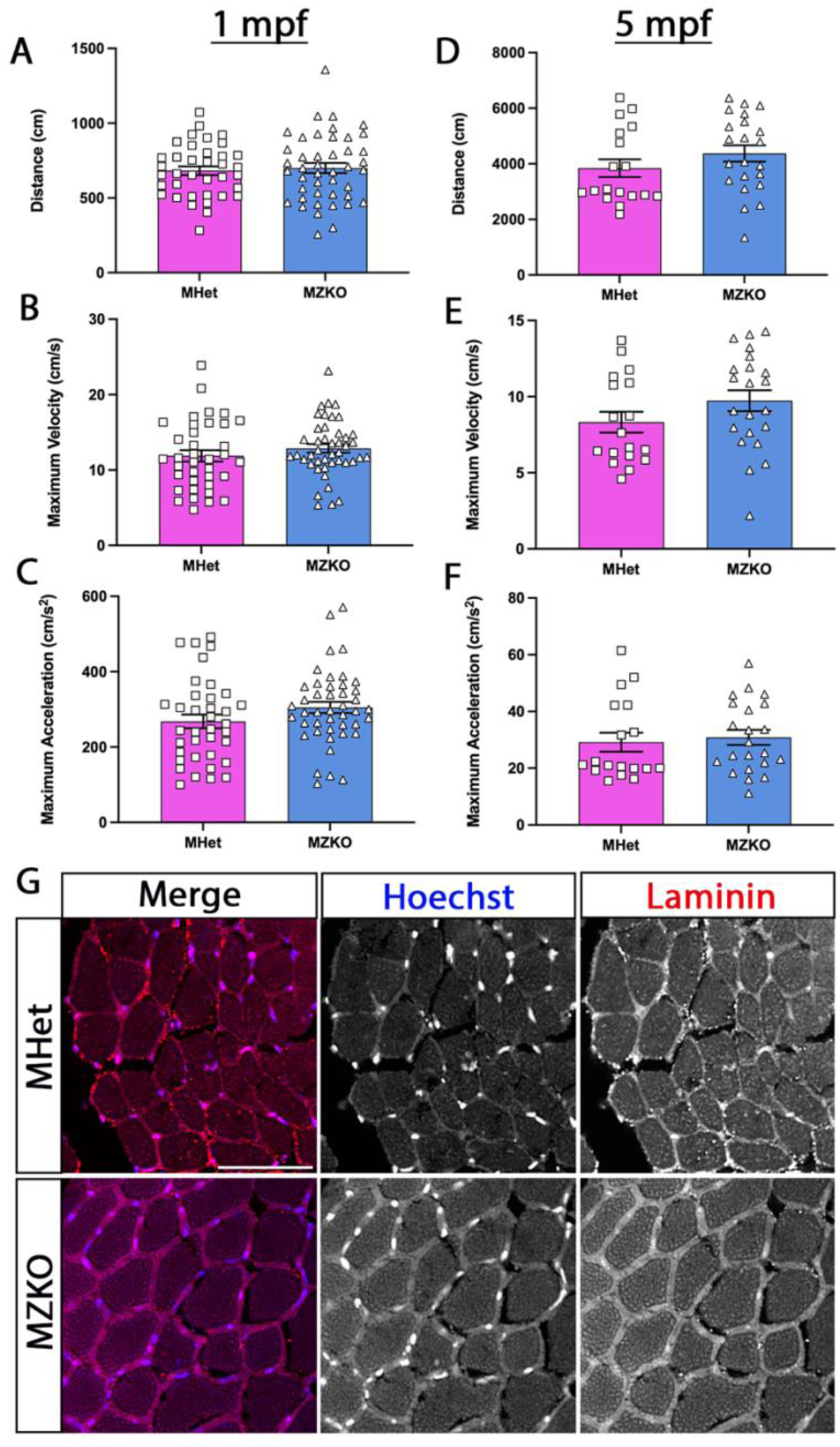
Behavioral and histological analysis does not indicate muscle disease in *pomgnt1* MZKO. **A-C**: Analysis of locomotor function at 1 mpf based on average distance swam, average maximum velocity, and maximum acceleration reveals no deficits in early development. Data are presented as mean ± SEM. **D-F**: Behavioral analysis at 5 mpf reveals no behavioral indications of muscle disease as zebrafish enter adulthood. Equal numbers of male and female zebrafish were tested. Data are presented as mean ± SEM. **G**. Images of 5 mpf transverse muscle sections stained with anti-laminin antibodies reveal no signs of muscle degradation or disorganization. Magnification: 20x. Scale bar: 50 µm.

Next, we assessed if any indicators of muscle disease were present on a histological level despite preserved locomotor function. Transverse muscle sections were taken at 5 mpf and immunostained for laminin to enable assessment of muscle fiber size and organization (**Fig.3G**). Muscle section integrity, organization, or muscle fiber size showed no differences between MHet and MZKO. These results taken with the locomotor assessments indicate no muscle disease in *pomgnt1*-deficient zebrafish at least up to 5 mpf.

While locomotor activity of 5 mpf zebrafish was tracked (**Fig. 4A**), we noticed that some fish displayed more wall-hugging behavior (thigmotaxis) and performed additional analyses of swim trajectories within the testing arena. The MZKO fish primarily swam along the perimeter of the arena, resulting in a significantly higher thigmotaxis score, whereas MHets explored the center to a higher degree (MHet: 0.787 ± 0.036 normalized thigmotaxis score, n=18; MZKO: 0.890 ± 0.010 normalized thigmotaxis score, n=22; N=2; p=0.0052) (**Fig. 4B**). Spatial entropy analysis was also used to assess differences in space utilization between the two groups, as the MZKOs spent significantly more time within a small portion of the perimeter while the MHets spent relatively equal amounts of time across all spatial bins of the testing arena (MHet: 0.884 ± 0.014, n=18; MZKO: 0.842 ± 0.007, n=22; N=2; p=0.0072) (**Fig. 4C-D**). Collectively, these results indicate that *pomgnt1* loss does not significantly affect gross locomotor activity but is associated with observable increases in thigmotaxis, a pattern often associated with anxiety-like behaviors in zebrafish (Srivastava et al., 2025).

**Figure 4.**
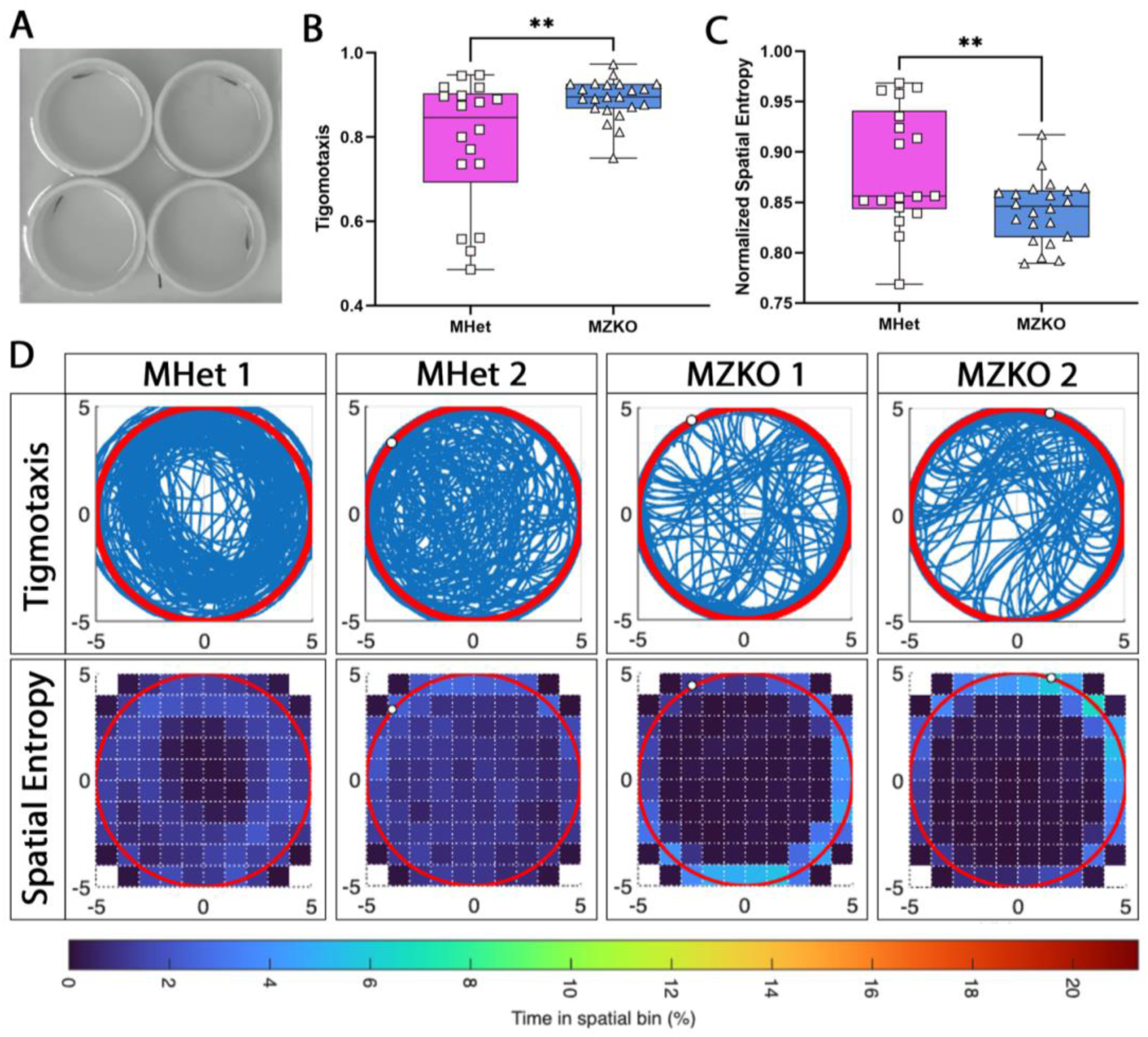
Abnormal swimming behavior in *pomgnt1* MZKO fish. **A.** Image of the individual testing arenas for the 5 mpf locomotor analysis experiments. **B**. Thigmotaxis scores indicate that MZKO fish spend significantly more time along the perimeter of the testing arena when compared to MHet counterparts. Data are presented as mean, minimum, and maximum values. **C**. Spatial entropy analysis shows that MZKO fish spend a significant portion of their time within a small section of the testing arena, while the MHet fish disperse their activity more evenly across the arena. Data are presented as mean, minimum, and maximum values. **D.** Tracing of individual swim trajectories of two MHets and two MZKOs are paired with heatmaps displaying their spatial entropy throughout the duration of the experiment, showing consistent overlap between the two parameters of analysis.

### Loss of maternal pomgnt1 reveals earlier retinal layering disorganization

To assess if retinal layering disruptions are seen earlier in MZKOs than in ZKOs, immunofluorescent staining was performed on head cryosections at 2 and 5 mpf. The cell bodies of photoreceptor cells in the outer nuclear layer (ONL) and their pedicles in the outer plexiform layer (OPL) were labeled using an anti-ZPR-1 antibody, which labels Arrestin 3, a critical transducer of light signals in photoreceptors (Renninger et al., 2011). An anti-synaptophysin (Syp) marker was used to visualize presynaptic vesicles in the OPL, where the photoreceptor cell pedicles synapse with bipolar and horizontal cells to support continuous transmission of visual signals.

At 2 mpf, MZKO fish exhibited consistent photoreceptor disorganization as well as sporadic protrusion of cone photoreceptor cell bodies into the OPL (**Fig. 5A-B**). By 5 mpf, a notable increase in the general disorganization of cell bodies was found in MZKO fish, with an increase in the number of protruding photoreceptor cell bodies into the OPL (**Fig. 5D-E**). In contrast, MHet retinae exhibited a consistently uniform localization of cone photoreceptor cell bodies at both timepoints. To quantify this trend, ZPR-1 signal intensity was measured within the portion of the ONL between the photoreceptor cell bodies and their pedicles and a statistically significant difference was found between the MHets and MZKOs at both timepoints (2 mpf: MHet: 14,101 ± 928 integrated signal density - IntDen, n=4; MZKO: 18,141 ± 1,050 IntDen, n=4; p=0.0279; 5 mpf: MHet: 14,641 ± 671 IntDen, n=5; MZKO: 20,319 ± 610 IntDen, n=4; p=0.0005) (**Fig. 5C, F**). While no significant differences in synaptophysin signal intensity were found, the distribution of the signal reflected the disorganization in the pedicles. These results demonstrate that *pomgnt1* loss disrupts photoreceptor organization within the ONL in a progressive manner, while synaptic organization is still preserved. These results identified a significant photoreceptor disorganization phenotype in MZKOs 4 months earlier than the mutant retinal phenotypes reported in ZKO progeny (Liu et al., 2020), indicating that maternally provided factors may delay the onset of retina disease pathology.

**Figure 5.**
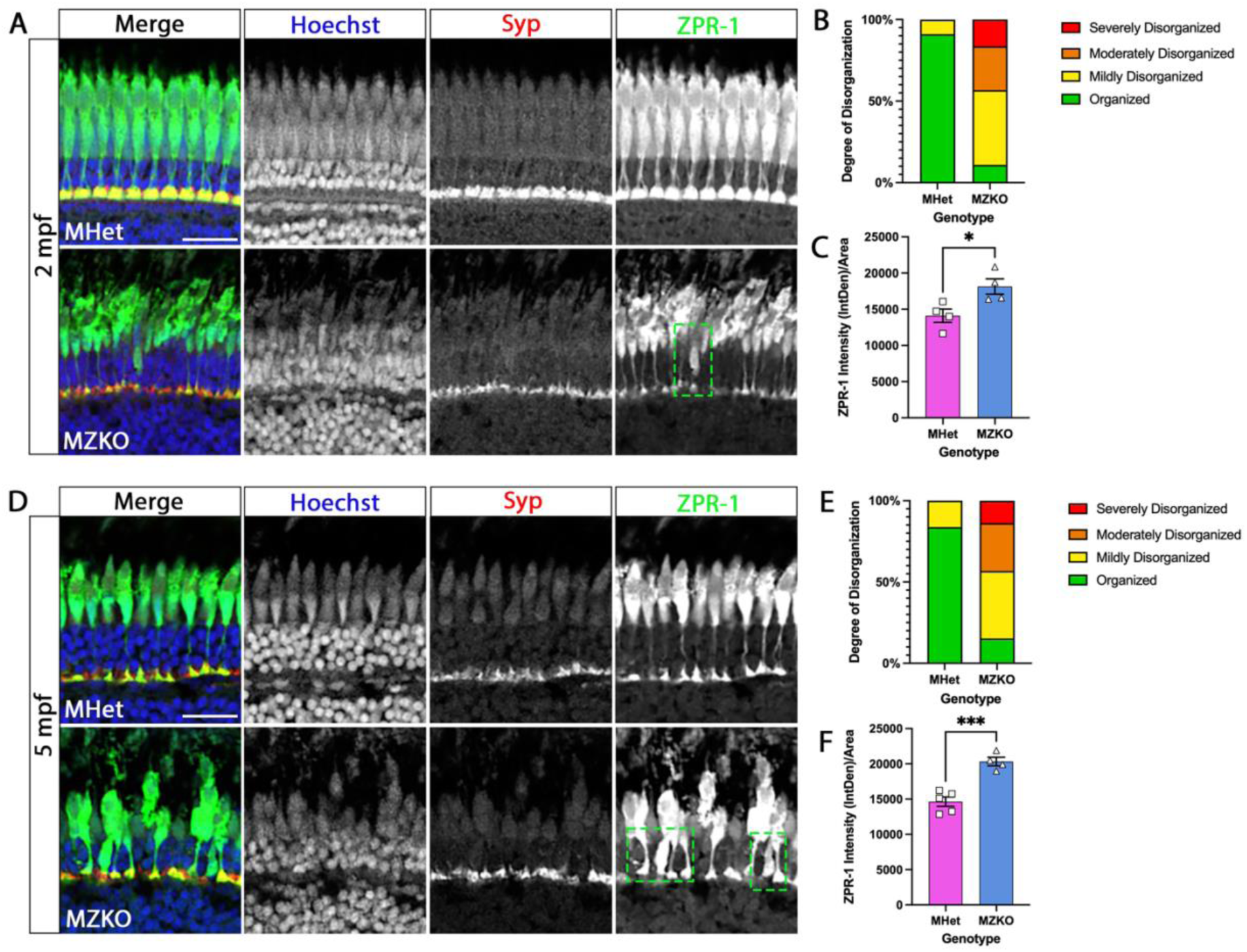
Analysis of retinal layering organization at 2 and 5 mpf. **A.** Comparison of immunofluorescent staining of the retina MHets and MZKOs at the 2 mpf timepoint shows the beginning disorganization of photoreceptor cell bodies in the ONL. Inlets show the OPL and ONL. Magnification: 20x. Scale bar: 20 µm. **B.** Stacked bar graph showing the varied distribution of the degree of disorganization seen across the two genotypes at 2 mpf**. C.** Quantification of displaced ZPR-1 signal intensity in portion of ONL between photoreceptor cell bodies and their pedicles shows significant difference between genotypes. Data are presented as mean ± SEM. **D.** Comparison of immunofluorescent staining of the retina MHets and MZKOs at the 5 mpf timepoint show continuation of disorganization trend seen at 2 mpf. Magnification: 20x. Scale bar: 20 µm. **E.** Stacked bar graph showing the varied distribution of the degree of disorganization across the two genotypes at 5 mpf. **F.** Quantification of displaced ZPR-1 signal intensity in portion of ONL between photoreceptor cell bodies and their pedicles shows significant difference between genotypes. Data are presented as mean ± SEM.

To further assess the severity of the retinal phenotype, we tested for localization and expression of cone-specific α-transducin using an anti-GNAT2 antibody. MZKO retinae exhibited a significant reduction in GNAT2 intensity in the outer segment of ZPR1-positive red/green double cone photoreceptors at the 2 mpf timepoint when compared to the MHets controls, accompanied by the same photoreceptor disorganization and cell body protrusions seen in the initial staining analyses (**Fig. 6A-B**. 2 mpf MHet: 1,962 ± 196 IntDen, N=4 fish; MZKO: 1,451 ± 59 IntDen, N=5 fish, p=0.0285). A statistically significant decrease in GNAT2 signal intensity in ZPR1-positive cells was also found at 5 mpf (5 mpf MHet: 1,682 ± 80 IntDen, N=4 fish; MZKO: 1,223 ± 101 IntDen, N=4 fish, p=0.0121) (**Fig. 6C-D**).

**Figure 6.**
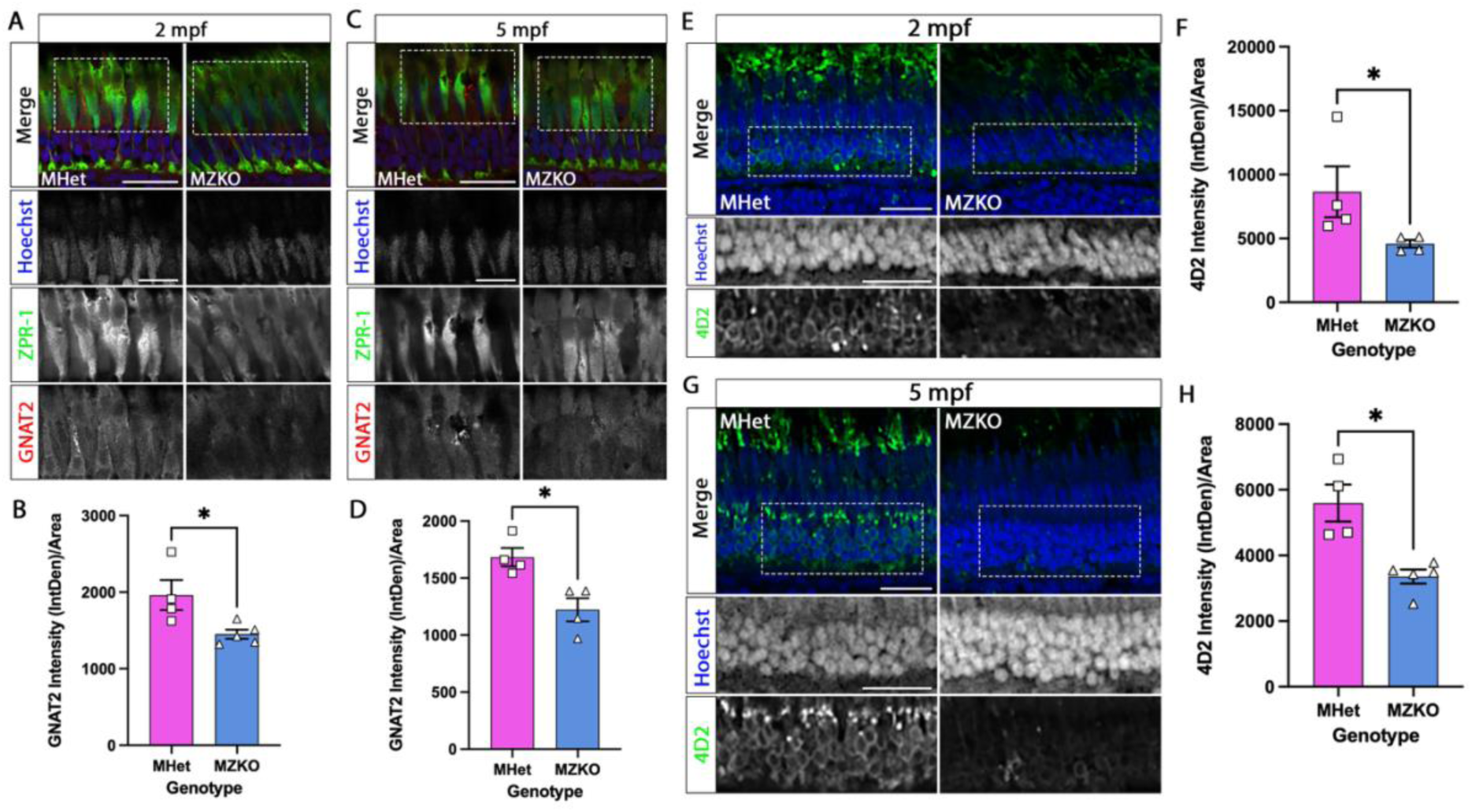
Disruptions in light transduction pathways in pomgnt1 MZKO fish. **A**: Fluorescent staining of retinal tissue from MHet and MZKO at 2 mpf shows reduced GNAT2 signal intensity in MZKOs. Insets focus on photoreceptor cell bodies. Magnification: 63x. Scale bar: 20 µm for merged image, 10 µm for greyscale insets. **B**: Quantification of GNAT2 signal intensity within cone photoreceptor cell bodies reveals significant reduction in MZKOs at 2 mpf. Data are presented as mean ± SEM. **C.** Fluorescent staining of retinal tissue from ZKOxHet progeny at 5 mpf shows expected reduction in GNAT2 signal intensity in MZKOs when compared to MHets. Magnification: 63x. Scale bar: 20 µm for merged image, 10 µm for greyscale insets. **D**: Quantification of GNAT2 signal intensity within cone photoreceptor cell bodies at 5 mpf confirms significant reduction in MZKOs. Data are presented as mean ± SEM. **E**. Fluorescent staining of the retina, with insets around the rod specific portion of the ONL. Results at 2 mpf show reduced GNAT2 signal intensity in the ONL-R of MZKOs when compared to MHets. Magnification: 20x. Scale bar: 20 µm for both the merged images and greyscale insets. **F**. Quantification of 4D2 signal intensity within ONL-R shows reduction in MZKOs at 2 mpf. Data are presented as mean ± SEM. **G**. Confocal images showing the 4D2 intensity difference between MHets and MZKOs at the 5 mpf timepoint, with insets of the ONL-R. Magnification: 20x. Scale bar: 20 µm for both the merged images and greyscale insets. **H**. 4D2 signal intensity analysis at 5 mpf shows an expected significant reduction of 4D2 signal in MZKOs compared to MHets. Data are presented as mean ± SEM.

In addition, we tested for rhodopsin expression using an anti-rhodopsin antibody (4D2) to visualize rod photoreceptors. At 2 mpf and 5 mpf, MZKO retinae exhibited a significant reduction in 4D2 signal intensity within the ONL relative to the MHets (**Fig. 6E,G**). The analysis revealed a significantly lower intensity in MZKOs than in the MHet controls at both 2 and 5 mpf (2 mpf: MHet: 8,652 ± 1,986 IntDen, N=4 fish; MZKO: 4,589 ± 298 IntDen, N=4 fish, p=0.0286; 5 mpf: MHet: 5,597 ± 561 IntDen, N=4 fish; MZKO: 3,356 ± 214 IntDen, N=5 fish, p=0.0159) (**Fig. 6F, H**). Collectively, these results demonstrate that both cone and rod associated light-conduction machinery is reduced at the 2 mpf timepoint in MZKOs. The same deficits were only found at 6 mpf in ZKO progeny and were not present at 2 mpf (Liu et al., 2020). The shift in timeline of disease phenotypes in the retina supports the earlier model where maternally provided factors buffer early retinal deficits in *pomgnt1* mutants.

## Discussion

In this study, we characterized a zebrafish maternal zygotic KO for the Pomgnt1 glycosyltransferase to determine whether this model could better capture phenotypic variability associated with *POMGNT1*-related disorders. Characterization of the retinal pathology of MZKO zebrafish revealed that maternal compensation delays the onset and severity of the retinal phenotypes previously reported in ZKOs, an important consideration for future studies using this model (Liu et al., 2020). Additionally, locomotor assessment and skeletal muscle histology revealed no overt muscle phenotype through 5 mpf, despite the association of *POMGNT1* loss with several forms of CMD in humans.

One possible explanation for the absence of the anticipated muscle phenotype is that evolutionary divergence between lower and higher vertebrates has resulted in species-specific differences in the dependence of skeletal muscle on POMGNT1-mediated glycosylation. In mammals, POMGNT1 may play a more critical role in supporting matriglycan maturation or have additional functions that have yet to be fully understood. Matriglycan itself could be sufficient to maintain skeletal muscle integrity in zebrafish, without the additional glycan maturation mediated by Pomgnt1. This possibility is supported by previous work with zebrafish models lacking *pomt1* or *pomgnt2*, two glycosyltransferase genes required for matriglycan initiation, both of which exhibit severe muscle disease pathology, locomotor deficits, and reduced viability (Flannery et al., 2026; Karas et al., 2024). In contrast, disruption of *pomgnt1*, which contributes to glycosylation of α-DG, but is not directly required for initial matriglycan formation, does not produce comparable muscle phenotypes in fish. These differences raise the possibility that the relationship between POMGNT1-mediated glycosylation and matriglycan function has diverged between humans and zebrafish, where matriglycan may be sufficient to maintain skeletal muscle integrity independently of POMGNT1-dependent maturation.

A second, non-mutually exclusive explanation for the absence of skeletal muscle deficits is that different tissues may have distinct requirements for functional *α*-DG glycosylation, with some tissues being more tolerant of reduced glycosylation while others requiring specific stages of glycan maturation. Previous work in mice has suggested the existence of a threshold of *α*-DG glycosylation where partial reductions are tolerated in skeletal muscle due to partial *α*-DG laminin-binding capacity (Kanagawa et al., 2009). In contrast, POMGNT1 has been shown to be critical for retinal layer organization and synapse formation in mice, and retinal ligands, such as pikachurin, depend on POMGNT1 activity for binding (Kanagawa et al., 2010). Retinal and brain *α*-DG shows less total glycosylation than muscle *α*-DG and binding may also depend on POMGNT1-extended glycans (Kanagawa et al., 2010; Sugita et al., 2001). Following loss of *pomgnt1*, the remaining *α*-DG glycosylation in zebrafish muscle could be sufficient to support tissue integrity, while loss of Pomgnt1 may have more severe functional consequences in the retina.

The disorganization of retinal layers and loss of photoreceptor cells observed in this study are consistent with those seen in *POMGNT1*-related RP in humans, which includes rod and cone loss accompanied by abnormal retinal layer organization (Ghanawi and Koch, 2025; Jones et al., 2016). This structural retinal pathology suggests that loss of *pomgnt1* produces retinal degeneration, and it could be used in studying *POMGNT1*-associated retinal disease. The focused use of this model is also supported by the limitations in modeling the neurological symptoms of brain malformations associated with CMD in zebrafish. Severe CMD presentations caused by loss of *α*-DG glycosylation often feature hydrocephalus, cortical malformations such as cobblestone lissencephaly and pachygyria resulting from abnormal neuronal migration, and severe intellectual impairment (Cormand et al., 2001; Haltia et al., 1997; Hehr et al., 2007). These deficits are orchestrated by several *α*-DG ligands involved in neuronal migration, axon guidance, and synaptic function (Jahncke and Wright, 2023). While retinal axon guidance deficits have been recapitulated in zebrafish (Flannery et al., 2026; Karas et al., 2024), to date, genetic models lacking *α*-DG glycosylation have not shown brain malformations (Flannery et al., 2026; Karas et al., 2024; Liu et al., 2022). As zebrafish lack a cerebral cortex and undergo different morphogenetic and neurogenetic processes (Jurisch-Yaksi et al., 2020), it is possible that these major structural differences and lack of certain ligand interactions prevent the development of brain malformations in fish. Dystroglycan (*dag1*) KO zebrafish die within 10 dpf, but brain development has not been carefully analyzed (Gupta et al., 2011). Additional work would be needed to determine whether synaptic deficits are present in the brain of zebrafish with disrupted *α*-DG function.

The behavioral abnormalities we report in 5 mpf *pomgnt1* MZKO zebrafish are intriguing as they could represent visual or central synaptic deficits. Thigmotaxis is consistent with phenotypes in Mexican blind cave fish (*Astyanax mexicanus*), a cave-adapted form of Mexican tetra that underwent eye loss (Ponnimbaduge Perera et al., 2023). Thigmotaxis in MZKO fish strongly resembles that of the “wall-hugging” behavior reported in these blind fish, which navigate through their environment using extrasensory inputs (Patton et al., 2010; Sharma et al., 2009). We propose that vision deficits contribute to behavioral changes in MZKOs. However, thigmotaxis has also been described as an anxiety-like phenotype in zebrafish models of neurodevelopmental disorders and could reflect additional synaptic dysfunction (Hiramoto et al., 2011).

The earlier onset of retinal degeneration in MZKOs indicates that maternal contribution substantially delays the manifestation of the retinal phenotype in ZKOs. Most maternal effects in zebrafish become apparent during gastrulation and early embryonic development (Solnica-Krezel, 2020). In zygotic mutants (ZKO) disrupting *α*-DG glycosylation deficits can appear well after a month post fertilization due to the long half-life of glycosylated *α*-DG in tissues (Flannery et al., 2026; Karas et al., 2024; Novak et al., 2021). However, when *α*-DG glycosylation initiation is disrupted by removal of *pomt1* or *pomgnt2*, developmental phenotypes are evident within a few days (Flannery et al., 2026; Karas et al., 2024). The appearance of retinal disease pathology at 2 mpf in MZKOs likely reflects a maintenance role for POMGNT1 activity in photoreceptor synapses, that should be considered when using this model to investigate POMGNT1-related disease mechanisms in the future.

The absence of overt muscle disease phenotypes accompanied by the robust retinal phenotype suggest that zebrafish *pomgnt1* loss produces a strong tissue-selective phenotype. This tissue selectivity provides an opportunity to investigate why some patients with POMGNT1 deficiency develop isolated retinal disease while others develop broader multisystem phenotypes. Given the strong retinal phenotype and the limited ability of this model to recapitulate brain malformations, this mutant line may be useful for understanding the mechanisms of POMGNT1-associated retinal degeneration and RP.

## Materials and Methods

### Mutant Lines and Animal Husbandry

The mutant *pomgnt1* line was generated via CRISPR/Cas9 genome editing by the Hu Lab at Upstate Medical University in Syracuse, NY (Liu et al., 2020). All experiments and procedures were conducted according to institutional guidance and approved by the Rutgers University Institutional Animal Care and Use Committee. Zebrafish were housed on a recirculating Tecniplast USA system under a 14-10 light-dark cycle at 28°C in 3.5 L tanks. For breeding events, adult zebrafish were placed off system in divided breeding tanks during the evening and dividers were removed the following morning at the beginning of the light cycle. Developing embryos were kept in Petri dishes and stored in an incubator at 28°C up to 5dpf and then moved onto the system at the standard stocking densities until the required timepoints. Crosses between heterozygous adults were used to generate WT and ZKO zebrafish and crosses between a female ZKO and male heterozygote produced MHets and MZKOs.

### Genotyping

For zebrafish older than 2 mpf, fin clipping was performed to determine genotype. Zebrafish were anesthetized using solution of 0.4% tricaine methanesulfonate (Tricaine, MS-222) solution in system water, then transferred to a clean surface to remove a small portion of their anal fin using a clean razor blade. For timepoints younger than 1 mpf, the fish were sacrificed and their tails were removed using a clean razor blade. Following DNA extraction using Extract-N-Amp Tissue PCR Kit (XNAT2-1KT, Millipore Sigma), genotyping was performed through quantitative allele-specific PCR with custom designed Affinity Plus™ qPCR probes and primers from Integrated DNA Technologies (Coralville, IA). These were run on a QuantStudio™ 6 qPCR system (Applied Biosystems) for 40 cycles.

### Morphological Analysis

To analyze survival, zebrafish of unknown genotypes were placed at maximum tank density to induce competition for food. The survival rates were determined by genotyping the remaining zebrafish at each timepoint and comparing the genotypic ratio of the survived fish against the expected Mendelian ratios. For most of these timepoints, the same fish were utilized to analyze body length. For the 1 mpf timepoint, the zebrafish were imaged in 3% methylcellulose with a M165 FC stereo microscope and LAS Software v4.21 (Leica Microsystems) using a ruler as a reference. For the 2 and 5 mpf timepoints, the fish were imaged with a handheld camera (iPhone 13) using a ruler as a reference. All images were then assessed in ImageJ by measuring body length from the mouth to the base of the tail fin. These images were assessed with the genotypes unknown to the experimenter to avoid bias.

### Automated Mobility Tracking

For behavior analysis at 1 mpf, animals were placed in individual wells of a 24 well polystyrene cell culture plate in a DanioVision™ Observation Chamber (Noldus Information Technology, Wageningen) and locomotor activity was tracked with EthoVision® XT Video Tracking software. Each well was filled with equal volumes of system water. Each plate was placed in the observation chamber for a total of 40 minutes (20 minute adaptation, 20 minute testing) at a temperature of 28.5°C. The fish were then fin clipped and utilized for other data points including survival, body morphology, and histology. All behavior experiments were performed at the same time of day (between 9am and 12pm) with the experimenter masked to genotype.

Locomotor function at 5 mpf was analyzed by placing zebrafish in individually in custom-designed, 3D-printed circular tanks (*D* = 10 cm in diameter) containing water maintained at approximately 26 ± 1.5°C. Each fish was allowed to habituate to the arena for 5 min, followed by a 10-min recording period using a GS3-U3-41C6NIR-C 1” FLIR Grasshopper video camera at 30 frames per second. Fish trajectories were extracted using AnimalTA (Chiara and Kim, 2023) and subsequently processed and analyzed in MATLAB. Tracking coordinates were converted from pixels to physical units, smoothed using a Kalman filter (Kalman, 1960), and centered relative to the arena prior to behavioral analysis. Thigmotaxis was quantified as the fraction of recording time spent in the peripheral region of the arena, defined as r ≥ 0.8R, where r is the radial distance from the arena center and R=D/2 is the arena radius. Spatial exploration was quantified using normalized Shannon entropy (Burbano-L. and Porfiri, 2021). The arena was divided into a 10 × 10 Cartesian grid, and the occupancy probability *p_j_* of each spatial bin was calculated from the proportion of recording time spent in that bin. Spatial entropy was computed as *H* = −*Σ_j_ p_j_ ln*(*p_j_*) (Shannon, 1948). To account for the finite number of spatial bins available within the circular arena, entropy was normalized by its theoretical maximum, *H*_max_ = ln(*N*_acc_), where is the number of grid cells intersecting the arena. Normalized spatial entropy was therefore defined as *H_norm_* = *H*/ ln(*N*_acc_), yielding values between 0 and 1. Higher values indicate more uniform spatial exploration, whereas lower values indicate more localized occupancy. All experiments were conducted at approximately the same time of day to minimize variability associated with circadian effects on ungenotyped fish using equal numbers of males and females. Following testing, fish were euthanized, fin-clipped, and measured for body morphology.

### Whole Mount Fluorescent Immunohistochemistry

Whole mount fluorescent immunohistochemistry was performed as previously described in (Flannery et al., 2026), using IIH6C4 primary antibody (AB_2617216, Hybridoma Bank, 1:50 dilution) and goat anti-mouse AlexaFluor 568 secondary antibody (A-11031, Thermo Fisher Scientific, 1:250 dilution).

### Cryosectioning and Fluorescent Immunohistochemistry

Zebrafish were sacrificed using 0.4% tricaine methanesulfonate (Tricaine, MS-222) solution. Fish of every age group (juvenile, and adult) were fixed in 4% paraformaldehyde (PFA) overnight at 4°C. To prepare the fish for cryomolds, they were first washed 3 times for 5 minutes in 1X phosphate buffered saline (PBS). Additionally, adult fish were decalcified for 1.5 hours in Cal-Ex solution (Fisher Scientific). The fish were then transferred to 15% sucrose with 0.2% sodium azide in 1X PBS for 1-2 days at 4°C, followed by additional incubation in 30% sucrose with 0.2% sodium azide in 1X PBS for another 1-2 days at 4°C. Zebrafish were then frozen in Tissue Freezing Medium on a mixture of isopentane and dry ice. Cryosectioning was performed on a Leica CM1850UV Cryostat (Leica Microsystems) at a thickness of 12 µm and captured on charged glass slides (VWRⓇ Superfrost Plus™ Slide, 48311-703). Slides were dried on a slide warmer at 37°C for 1 hour and either stained immediately or stored at −20°C.

For all histology experiments, sections were blocked in 10% normal goat serum (NGS) with 0.1% Triton X-100 (Tx100) for 1 hour. The primary staining was performed using 1% NGS with 0.1% Tx100 and the following primary antibodies: ZPR-1 (DSHB, 1:100), anti-synaptophysin (ab32127, AbCam, 1:300), GNAT2 (A10352, Abclonal, 1:100), and anti-rhodopsin (4D2, ab98887, Abcam, 1:300), all diluted in 1X PBS. These were incubated overnight at 4°C in a humidified chamber. On the following day the slides were washed 3 times for 5 minutes in 1X PBS followed by a secondary antibody incubation using 1%NGS with 0.1% Tx100 using Alexa 568 anti-rabbit IgG antibody and Alexa 488 anti-mouse IgG antibody (all at 1:500) for one hour at 4°C in a humidified chamber. Slides were before washed 3 times for 5 minutes in 1X PBS adding Hoechst 33342 (Thermo Fisher, 1:1000) as a nuclear stain. The slides were then mounted for imaging using ProLong™ Gold Antifade Mountant (Thermo Scientific, P36934). Slides were imaged on the Zeiss LSM800 confocal microscope using Zeiss Plan-APOCHROMAT objective: air immersion 10X (NA=0.45) and 20X (NA=0.8), and oil-immersion 40X (NA=1.4) and 63X (NA=1.4).

### Image analysis

Fiji/ImageJ was utilized for signal intensity quantification of confocal images (Schindelin et al., 2012). The ZPR-1/synaptophysin and 4D2 slides were imaged at 20X and the GNAT2/ZPR-1 slides were imaged at 63X. For the 4D2 signal, an ROI of 100 μm in length was drawn along the retina to capture the rod-specific portion of the ONL (ONL-R) for each image. Similarly, GNAT2 intensity analysis was performed by defining an ROI of 100 μm in length along the cone-specific portion of the ONL (ONL-C), starting from the bottom of the photoreceptor cell bodies and using the ZPR-1 stain as a guide. For the synaptophysin/ZPR-1 signal, ROIs spanned the lower border of the photoreceptor cell bodies and the outer plexiform layer. The integrated signal density and area were recorded for all ROIs, as well as the integrated signal density from non-stained areas of each image to account for background noise. To normalize any differences in area, the integrated density for each image was divided by the ROI area. This process was performed on 18-24 retinal image sections from each fish, the results being averaged and used for statistical analysis.

For the quantitative assessment of photoreceptor cell body disorganization in the ZPR-1/synaptophysin stains, each image was visually scored for the number of cell body protrusions and gaps within the ONL. Organized categorization referred to neat, tightly packed, parallel photoreceptor cell bodies with 0 protrusions. Mildly disorganized was characterized by 1-2 mild protrusions. Moderately disorganized referred to 3-5 protrusions or instances of reduced parallel packing of photoreceptor cell bodies. Severely disorganized refers to 6+ protrusions and/or large, distinct gaps in the ONL photoreceptor cell bodies. 18-20 retinal images were analyzed per fish at set retinal locations and the results averaged.

### Statistical Analysis

All statistical analyses were performed using GraphPad Prism v.8.20 (GraphPad, San Diego, CA). Data were assessed for normality and analyzed with appropriate parametric (Student t-test) or non-parametric (Mann-Whitney test) tests. Statistical significance was defined as a p-value of <0.05, with degrees of significance as follows: (*<0.05; **<0.01; ***<0.001; ****<0.0001). Each datapoint was representative of an individual fish (n), except for the survival analyses. For survival analyses, each datapoint represents the percentage of each genotype within an independent clutch of fish (N). The total number of individual fish (n) per clutch is included in the figure legend for reference. All error bars represent the standard error of the mean (SEM).

## Acknowledgements

We would like to thank Kathleen Flaherty of the Rutgers Zebrafish Facility and Rutgers Animal Care staff members for her incredible dedication and care in maintaining the zebrafish core facility. We would also like to thank Dr. Huaiyu Hu at Upstate Medical University for providing the *pomgnt1* zebrafish line used in this study.

## Author Contributions

Conceptualization: M.C.M., C.O.; Formal analysis: C.O., M.C.M., D.L., D.B., K.F.; Funding acquisition: M.C.M.; Investigation: C.O., K.F., L. S., L. S. D.L.; Visualization: C.O.; Project administration: C.O.; Supervision: M.C.M.; Writing – original draft: C.O.; Writing – review C editing: M.C.M., C.O.

